# Prolactin links psychological stress to psoriasis via a fibroblast-chemokine pathway

**DOI:** 10.1101/2025.10.11.681540

**Authors:** Huiyao Ge, Shiyao Zhou, Yu Zhao, Xinwen Zhang, Mei Chen, Pan Gao, Shengxiu Liu, Qitian Mu, Difeng Fang, Xin Hou, Liang Yong

## Abstract

Psoriasis is a chronic inflammatory skin disease that forms a vicious cycle with psychological stress. Whether and how the hypothalamic-pituitary (HP) axis-mediated neuroendocrine system regulates psoriasis remains obscure. Here, we report elevated levels of the pituitary hormone prolactin (PRL) in both psoriasis patients and imiquimod (IMQ)-induced psoriasis mice. Mechanistically, PRL acts on dermal PRLR-expressing fibroblasts to promote the production of the chemokines CCL2 and CCL7, which then recruit monocytes/macrophages into psoriatic lesional skin, thereby activating local IL-17A-producing T cells. Accordingly, pharmacological targeting of PRL signaling inhibits the recruitment of monocytes/macrophages, decreases the frequency of IL-17A-producing T cells, and alleviates IMQ-induced psoriasis in mice. In summary, our results delineate a mechanism by which the neuroendocrine hormone PRL aggravates psoriasis and highlight a potential therapeutic strategy of inhibiting PRL-PRLR signaling, particularly in psoriasis patients experiencing psychological stress.

## Introduction

Psoriasis is an immune-mediated, chronic, and inflammatory skin disease associated with environmental and genetic factors, which affects approximately 125 million people worldwide^1,2^. Psoriasis not only causes physical symptoms, such as erythema, scales, itching, and pain, but also may lead to serious psychological stress and mental illnesses^3,4^. This stress leads to a worsening of the disease, establishing a deleterious feedback loop that adversely affects psoriasis prognosis and reduces overall quality of life^3,5,6^. Stress is recognized as a significant contributing factor in the onset and exacerbation of psoriasis^2,7^.

Psychological stress leads to the activation of the central nervous system (CNS)^8^. Both epidemiologic and experimental studies both have shown that the CNS regulates local skin immune responses and participates in the pathogenesis of psoriasis^9,10^. Either surgical denervation or lidocaine treatment results in a significant improvement in acanthosis and dermatitis in a mouse model of psoriasis^11,12^. Neurotransmitters derived from skin sensory nerves, such as substance P and calcitonin gene-related peptide, activate dendritic cells (DCs) to drive immune responses^11-13^. Additionally, nociceptive sensory neurons directly interact with DCs and γδT cells to promote psoriasiform dermatitis^14,15^. The neuroendocrine system, mediated by the HP axis, is another pathway of the CNS that can regulate the responses of downstream target organs^16^. The role of pituitary-derived hormones in regulating psoriasis remains unclear.

Skin fibroblasts, resided in the dermis, can be categorized into papillary, reticular, and hair follicle-associated subsets^17^. In addition to differences in anatomical location and morphology, dermal fibroblasts exhibit functional heterogeneity in skin homeostasis and disease^17,18^. Based on their functions in the processes of skin inflammation, immunity, and repair, dermal fibroblasts can be further classified into distinct functional subsets^15,19,20^. Emerging evidence from single-cell RNA sequencing indicates that various pathogenic fibroblast subsets, such as WNT5A^+^, SFRP2^+^, COL6A5^+^, and TNC^+^ fibroblasts, are involved in the pathogenesis of psoriasis by secreting cytokines and chemokines to regulate immune cells^15,21-23^.

Here, we identify a unique PRLR^+^ fibroblast subset that links the neuroendocrine system with psoriasis. Mechanistically, PRLR^+^ fibroblasts that receive PRL signaling secrete CCL2 and CCL7 to recruit monocytes/macrophages into psoriatic lesional skin, subsequently activating local IL-17A-producing T cells through release IL-1β and IL-23. Importantly, small molecule- and neutralizing antibody-mediated inhibition of PRL significantly improves psoriasis in mice. Our findings provide insight into the mechanism by which the neuroendocrine hormone PRL aggravates psoriasis, with implications for therapeutic approaches aimed at blocking PRL-PRLR signaling in psoriasis patients with psychological stress.

## Results

### PRL bridges psoriasis and stress

To investigate the relationships between psoriasis and psychological factors, we recorded psoriasis area and severity index (PASI), anxiety score GAD-7, and depression score PHQ-9 of psoriasis patients and analyzed their correlations. We found that PASI was positively correlated with GAD-7 and PHQ-9 (**Fig. 1a, b**). Similarly, we found that IMQ-induced psoriasis mice exhibited anxiety- and depression-like behaviors by the elevated plus maze (EPM) and open-field test (OFT) (**Fig. 1c, d**). These data indicate that psoriasis correlates with stress.

**Fig. 1.**
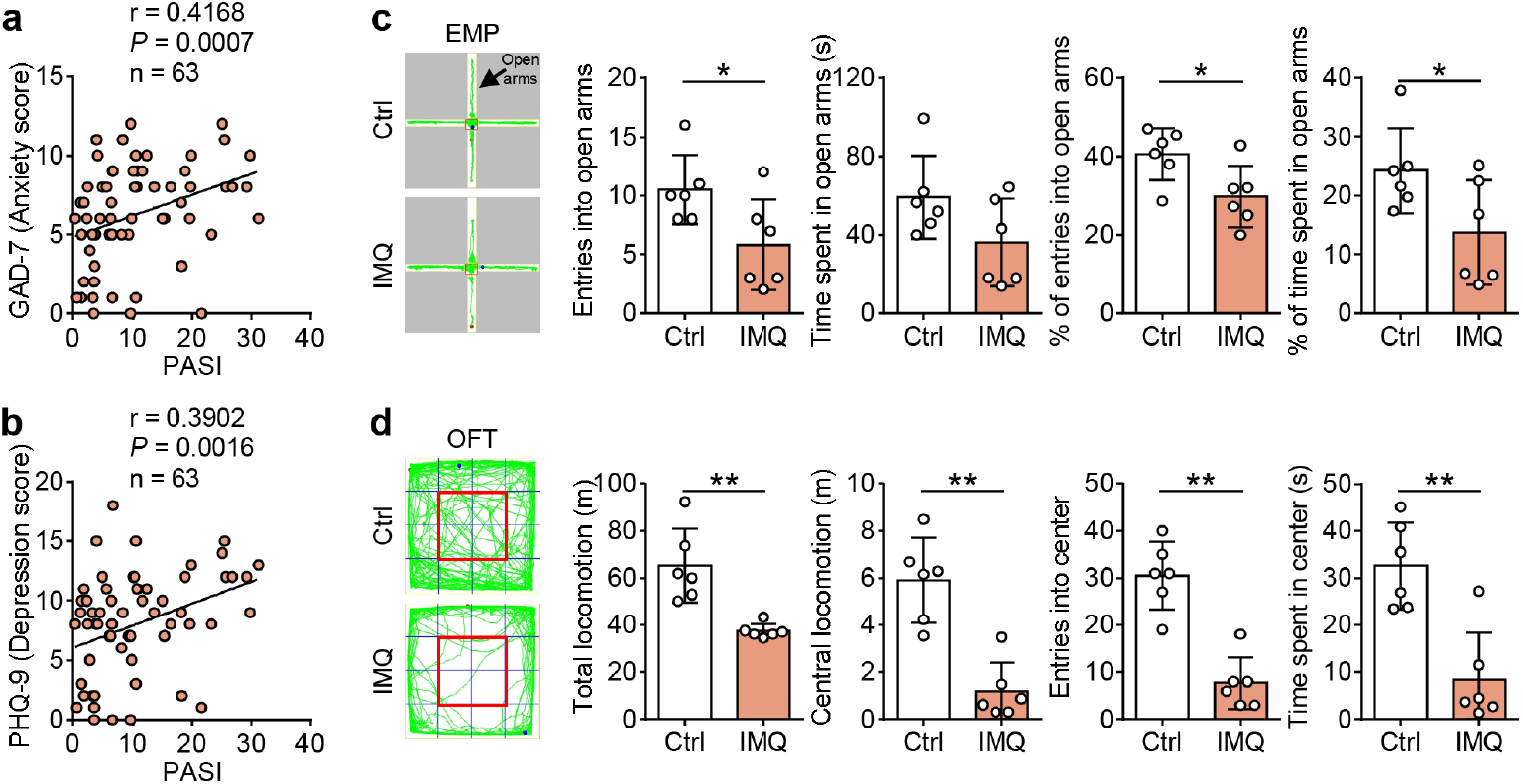
Psoriasis correlates with stress. **a, b** Correlations of PASI with GAD-7 (**a**) and PHQ-9 (**b**) in psoriasis patients (*n* = 63). **c, d** Representative locomotion tracks (green lines) of control (Ctrl) and IMQ mice (*n* = 6) in the EPM (**c**) and OFT (**d**). The red box indicates the central zone. Entries into open arms, time spent in open arms, the percentage of entries into open arms, and the percentage of time spent in open arms of the EPM (**c**) and total locomotion, central locomotion, entries into center, and time spent in center of the OFT (**d**) were compared between the two groups. Data are from two (**c, d**) independent experiments. Data are mean ± SD (**c, d**). The *P* values were calculated by Spearman’s rank correlation test (**a, b**) or two-tailed unpaired Student’s *t*-test (**c, d**); \**P* < 0.05, \*\**P* < 0.01.

We next sought clues linking psoriasis and stress within the HP axis. We examined pituitary mRNA levels and serum protein levels of pituitary-secreted hormones in IMQ-induced psoriasis mice. We found that pituitary mRNA levels of *Prl, Tshb*, and *Pomc*, and serum protein levels of PRL, TSH, LH, and ACTH were significantly increased in IMQ-induced psoriasis mice compared to the control mice (**Fig. 2a, b and Supplementary Fig. 1**), suggesting that the HP axis is activated in psoriasis.

**Fig. 2.**
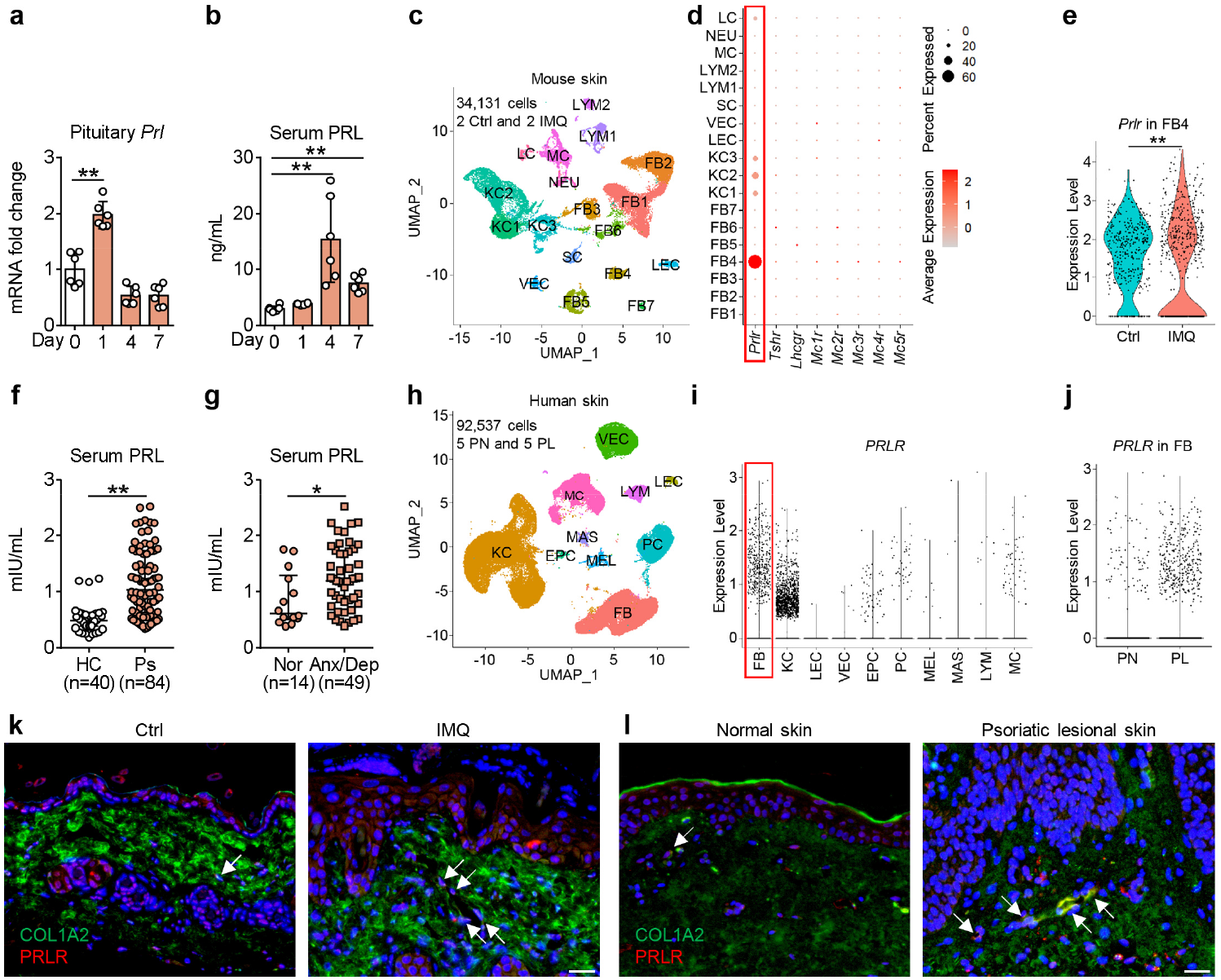
PRL is increased in psoriasis and its receptor PRLR is specifically and highly expressed in dermal fibroblasts. **a, b** Pituitary mRNA levels (**a**) and serum protein levels (**b**) of PRL in IMQ-treated mice at the indicated timepoints (*n* = 6). **c** UMAP plot of mouse skin cells from GSE165021. **d** Dot plot showing the expression of hormone receptors in different types of cells. **e** Violin plot of *Prlr* expression in the FB4 fibroblast subset. **f, g** Serum PRL levels of healthy controls (HC, *n* = 40) and psoriasis patients (Ps, *n* = 84) (**f**), as well as psoriasis patients with normal (Nor, *n* = 14) and anxiety and/or depression (Anx/Dep, *n* = 49) conditions (**g**). **h** UMAP plot of human skin cells from GSE228421. **i** Violin plot showing the expression of *PRLR* in different types of cells. **j** Violin plot of *PRLR* expression in FB. **k, l** Representative immunofluorescence images of skin sections stained with anti-COL1A2 (green), anti-PRL (red), and DAPI (blue) in mouse (*n* = 6) (**k**) and human (*n* = 4) samples (**l**). The solid arrows show PRLR^+^ fibroblasts. Scale bar, 20 μm. Data are from two (**a, b**) independent experiments. Data are mean ± SD (**a, b**) or median (IQR) (**f, g**). The *P* values were calculated by one-way ANOVA with Dunnett’s post-hoc test (**a, b**) or Mann-Whitney U test (**e**–**g**); \**P* < 0.05, \*\**P* < 0.01.

Based on the principle that the biological function of hormones is mediated by their receptors, we analyzed the expression of hormone receptors in different types of cells from mouse psoriatic lesional skin using a single-cell RNA sequencing dataset from public database. We observed that, among the hormone receptors, only *Prlr* was expressed in skin cells, and it was highly expressed in fibroblasts, whereas weakly expressed in keratinocytes (KCs) (**Fig. 2c, d and Supplementary Fig. 2a, b**). Moreover, the expression of *Prlr* was higher in fibroblasts of IMQ-induced psoriasis mice than in those of the control mice (**Fig. 2e**). Meanwhile, serum PRL levels were elevated in psoriasis patients compared to healthy controls (**Fig. 2f**). Additionally, serum PRL levels were higher in psoriasis patients with anxiety and/or depression than in those without these conditions (**Fig. 2g**). We also observed high expression of *PRLR* in fibroblasts and low levels of *PRLR* in KCs using a single-cell sequencing dataset of human psoriatic lesional skin (**Fig. 2h-j and Supplementary Fig. 2c, d**). We confirmed the expression of PRLR in fibroblasts and KCs through immunofluorescence staining both in mouse and human skin (**Fig. 2k, l and Supplementary Fig. 3a, b**). Taken together, these data demonstrate that the pituitary hormone PRL bridges psoriasis and stress, possibly via PRLR in fibroblasts or KCs.

### Deletion of *Prlr* in fibroblasts alleviates IMQ-induced psoriasis and stress-aggravated psoriasis

To investigate whether PRL participates in the pathogenesis of psoriasis via PRLR in fibroblasts or KCs, we generated *Prlr*^fl/fl^ *Col1a2*^Cre/wt^ and *Prlr*^fl/fl^ *KRT14*^Cre/wt^ mice in which *Prlr* was ablated in fibroblasts and KCs, respectively. The severity of psoriasis was alleviated in IMQ-treated *Prlr*^fl/fl^ *Col1a2*^Cre/wt^ mice compared to IMQ-treated *Prlr*^fl/fl^ mice (**Fig. 3a-c**). However, there were no difference in the histological phenotype of psoriasis between *Prlr*^fl/fl^ *KRT14*^Cre/wt^ and *KRT14*^Cre/wt^ mice (**Supplementary Fig. 3c-e**). These data indicate that PRL contributes to the pathogenesis of psoriasis via PRLR in fibroblasts.

**Fig. 3.**
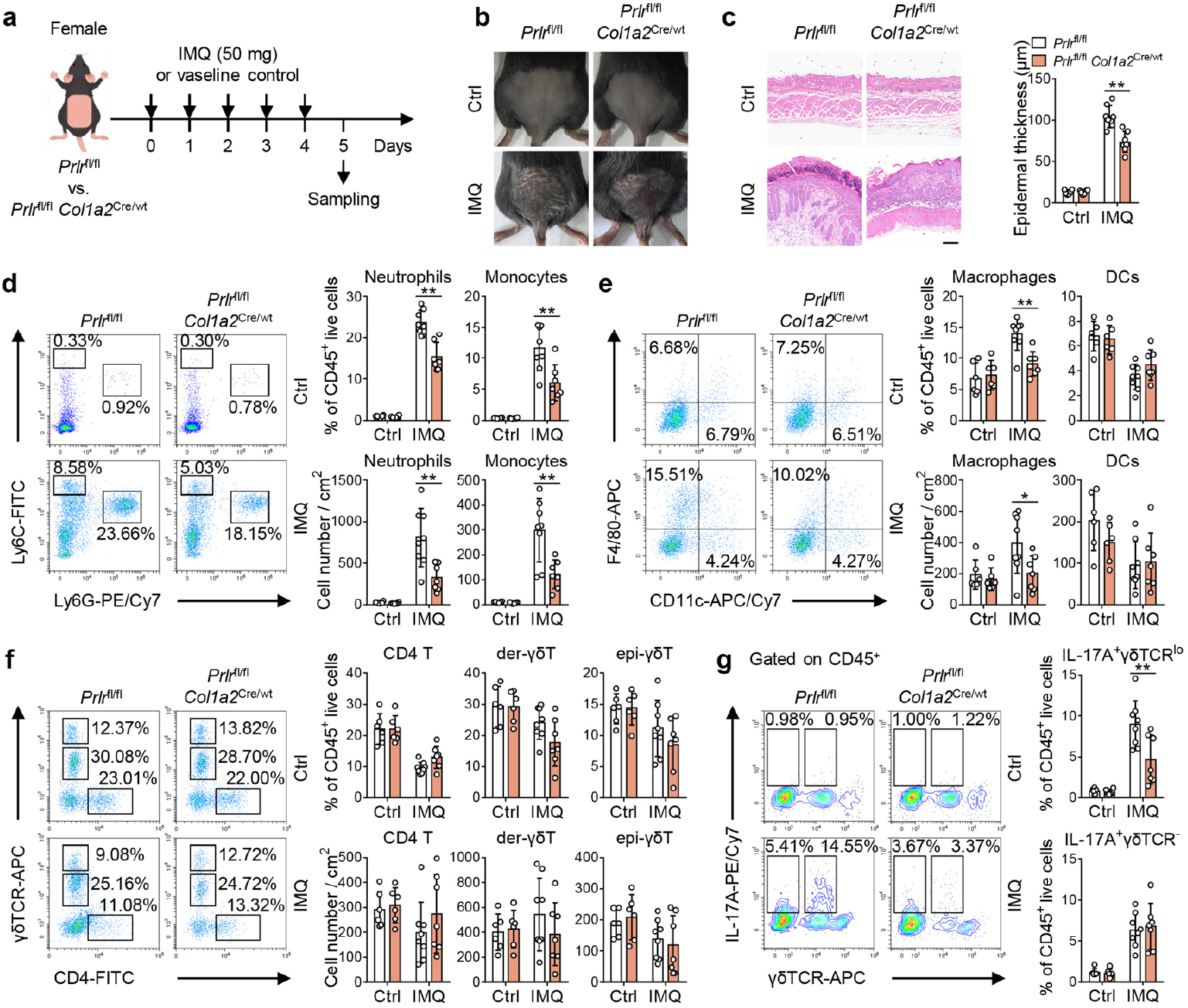
Deletion of *Prlr* in fibroblasts alleviates IMQ-induced psoriasis in female mice. **a** Schematic representation of the mouse model. **b** Representative photos of mouse back skin. **c** Representative H&E staining images and statistical analysis of epidermal thickness in Ctrl (*n* = 6) and IMQ (*n* = 7–8) mice. Scale bar, 100 μm. **d**–**g** Representative flow cytometry plots and statistical analyses of neutrophils and monocytes (**d**), macrophages and DCs (**e**), CD4 T, der-γδT, and epi-γδT cells (**f**), and IL-17A^+^ γδTCR^lo^ and IL-17A^+^ γδTCR^-^ cells (**g**) in the skin of Ctrl (*n* = 6) and IMQ (*n* = 7–8) mice. Numbers in representative plots indicate the frequency of each cell subset among CD45^+^ cells. Data are from two independent experiments. Data are mean ± SD (**c**–**g**). The *P* values were calculated by two-way ANOVA with Tukey’s post-hoc test (**c**–**g**). \**P* < 0.05, \*\**P* < 0.01.

Next, we analyzed local skin immune microenvironment of *Prlr*^fl/fl^ *Col1a2*^Cre/wt^ and *Prlr*^fl/fl^ mice using flow cytometry. The percentages and numbers of neutrophils, monocytes, and macrophages were markedly decreased in the skin of IMQ-treated *Prlr*^fl/fl^ *Col1a2*^Cre/wt^ mice compared to IMQ-treated *Prlr*^fl/fl^ mice, and there were no difference in DCs, CD4 T, der-γδT, and epi-γδT cells between the two groups of mice (**Fig. 3d-f and Supplementary Fig. 4**). Furthermore, the percentage of IL-17A^+^ γδTCR^lo^ cells was lower in the skin of IMQ-treated *Prlr*^fl/fl^ *Col1a2*^Cre/wt^ mice than in those of IMQ-treated *Prlr*^fl/fl^ mice (**Fig. 3g**).

To evaluate the influence of gender, we compared serum PRL levels between females and males in both mice and humans. In mice, serum PRL levels were comparable between females and males in both the control and IMQ-treated groups. Similarly, in humans, no significant difference was found between females and males among either healthy controls or psoriasis patients. (**Supplementary Fig. 5**). Similar to female mice, IMQ-treated *Prlr*^fl/fl^ *Col1a2*^Cre/wt^ male mice displayed less severe IMQ-induced psoriasiform dermatitis and reduced neutrophils, macrophages, and IL-17A^+^ γδTCR^lo^ cells in the skin relative to IMQ-treated *Prlr*^fl/fl^ male mice (**Supplementary Fig. 6**). Accordingly, the expression of *Tnf, Il1b, Il17a*, and *Il23a* were downregulated in IMQ-induced lesional skin of both female and male *Prlr*^fl/fl^ *Col1a2*^Cre/wt^ mice compared with those in *Prlr*^fl/fl^ mice (**Supplementary Fig. 7**). Collectively, these data suggest that PRL acts on fibroblasts and indirectly affects local skin immune responses to promote psoriasiform dermatitis.

Moreover, PRL supply exacerbated IMQ-induced psoriasiform dermatitis in *Prlr*^fl/fl^ mice, which was abolished in *Prlr*^fl/fl^ *Col1a2*^Cre/wt^ mice (**Fig. 4a, b**). This result further confirms that PRL promotes psoriasis via PRLR in fibroblasts.

**Fig. 4.**
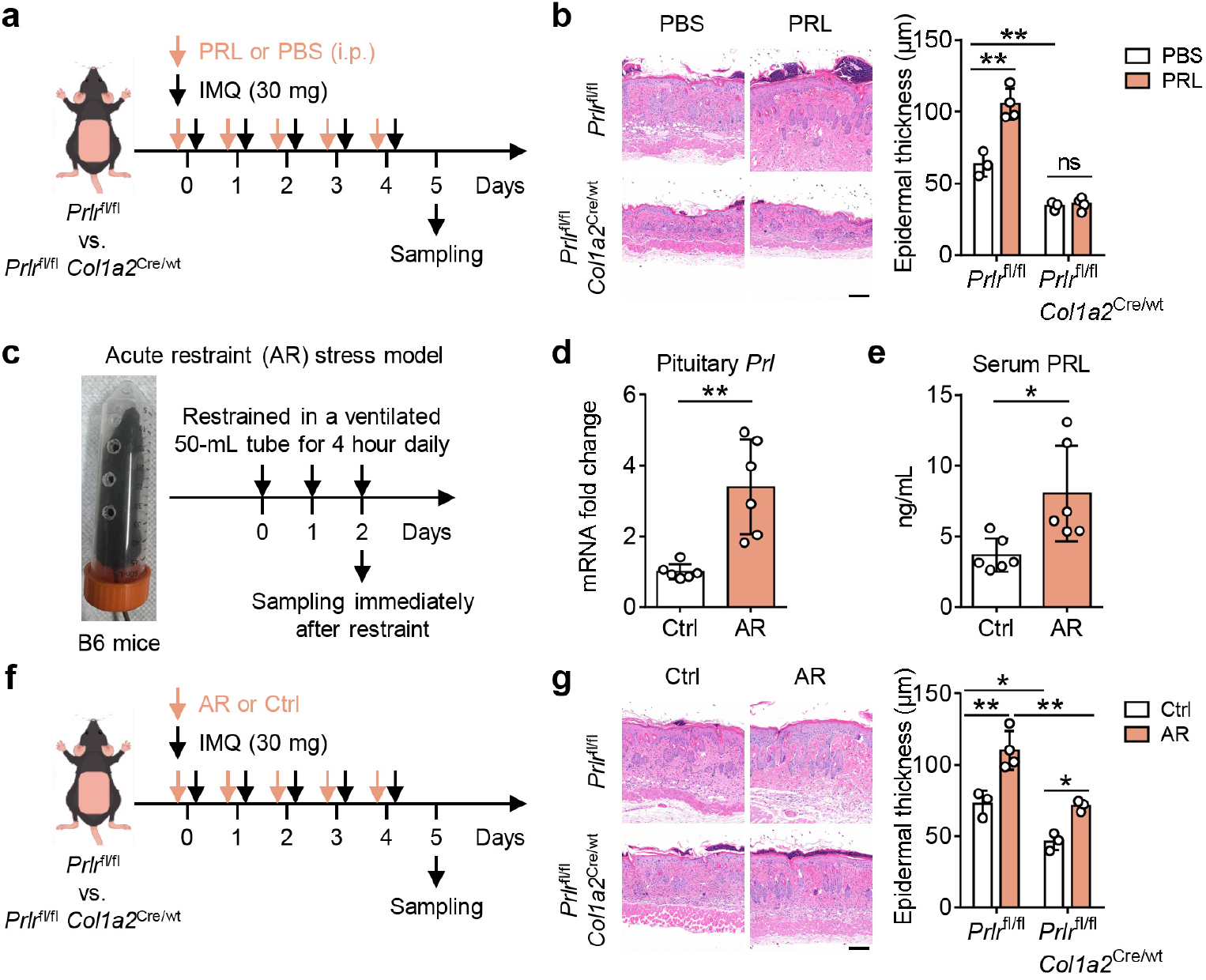
Ablation of *Prlr* in fibroblasts partially improves AR-aggravated psoriasis. **a** Schematic representation of the mouse model. **b** Representative H&E staining images and statistical analysis of epidermal thickness in PBS and PRL mice (*n* = 3–4). Scale bar, 100 μm. **c** Schematic representation of the mouse model. **d, e** Pituitary mRNA levels (**d**) and serum protein levels (**e**) of PRL in Ctrl and AR mice (*n* = 6). **f** Schematic representation of the mouse model. **g** Representative H&E staining images and statistical analysis of epidermal thickness in Ctrl and AR mice (*n* = 3–4). Scale bar, 100 μm. Data are from one (**a, b, f, g**) or two (**c**–**e**) independent experiments. Data are mean ± SD (**b, d, e, g**). The *P* values were calculated by two-way ANOVA with Tukey’s post-hoc test (**b, g**) or two-tailed unpaired Welch’s *t*-test (**d, e**); \**P* < 0.05, \*\**P* < 0.01, ns, not significant.

We then investigated whether stress aggravates psoriasis through the PRL-PRLR^+^ fibroblasts axis. Acute restraint (AR)-induced stress is a common approach to mimic psychological stress in mice^24^. AR could induce an increase in pituitary mRNA levels and serum protein levels of PRL and exacerbate the severity of IMQ-induced psoriasis. Ablation of *Prlr* in fibroblasts partially ameliorated AR-aggravated psoriasis (**Fig. 4c-g**), providing direct experimental evidence for stress exacerbating psoriasis through the PRL-PRLR^+^ fibroblasts axis.

### PRL promotes the recruitment of monocytes/macrophages through the JAK2-STAT3-CCL2/CCL7 axis in fibroblasts

To further explore the molecular mechanism of how PRL-PRLR signaling regulates fibroblasts, we stimulated fibroblasts with PRL and performed RNA sequencing (**Fig. 5a**). The chemokine signaling pathway was significantly enriched in PRL-stimulated NIH-3T3 cells through KEGG pathway enrichment analysis and GSEA (**Fig. 5b-e**). Additionally, the prolactin signaling pathway was also enriched (**Fig. 5c**). We used quantitative PCR to verify that the expression of *Ccl2* and *Ccl7* was significantly upregulated in NIH-3T3 cells stimulated by PRL compared to the control group (**Fig. 5f**). Consistently, the expression of *Ccl2* and *Ccl7* was downregulated in the skin of IMQ-treated female and male *Prlr*^fl/fl^ *Col1a2*^Cre/wt^ mice compared with that in *Prlr*^fl/fl^ mice (**Fig. 5g**). We also found that the JAK-STAT signaling pathway was enriched in NIH-3T3 cells upon PRL stimulation (**Fig. 5h**), which is consistent with existing reports that JAK-STAT signaling is downstream of the PRL-PRLR axis^25^. PRL stimulation induced the activation of JAK2 and STAT3, rather than STAT1 and STAT5 (**Fig. 5i**). These activations were inhibited in *Prlr* knockdown cells (**Fig. 5j**). Importantly, PRL-induced *Ccl2* and *Ccl7* expression was suppressed by a JAK2/STAT3 inhibitor (**Fig. 5k**). These data demonstrate that PRL-PRLR signaling promotes the production of CCL2 and CCL7 in fibroblasts via the JAK2-STAT3 signaling.

**Fig. 5.**
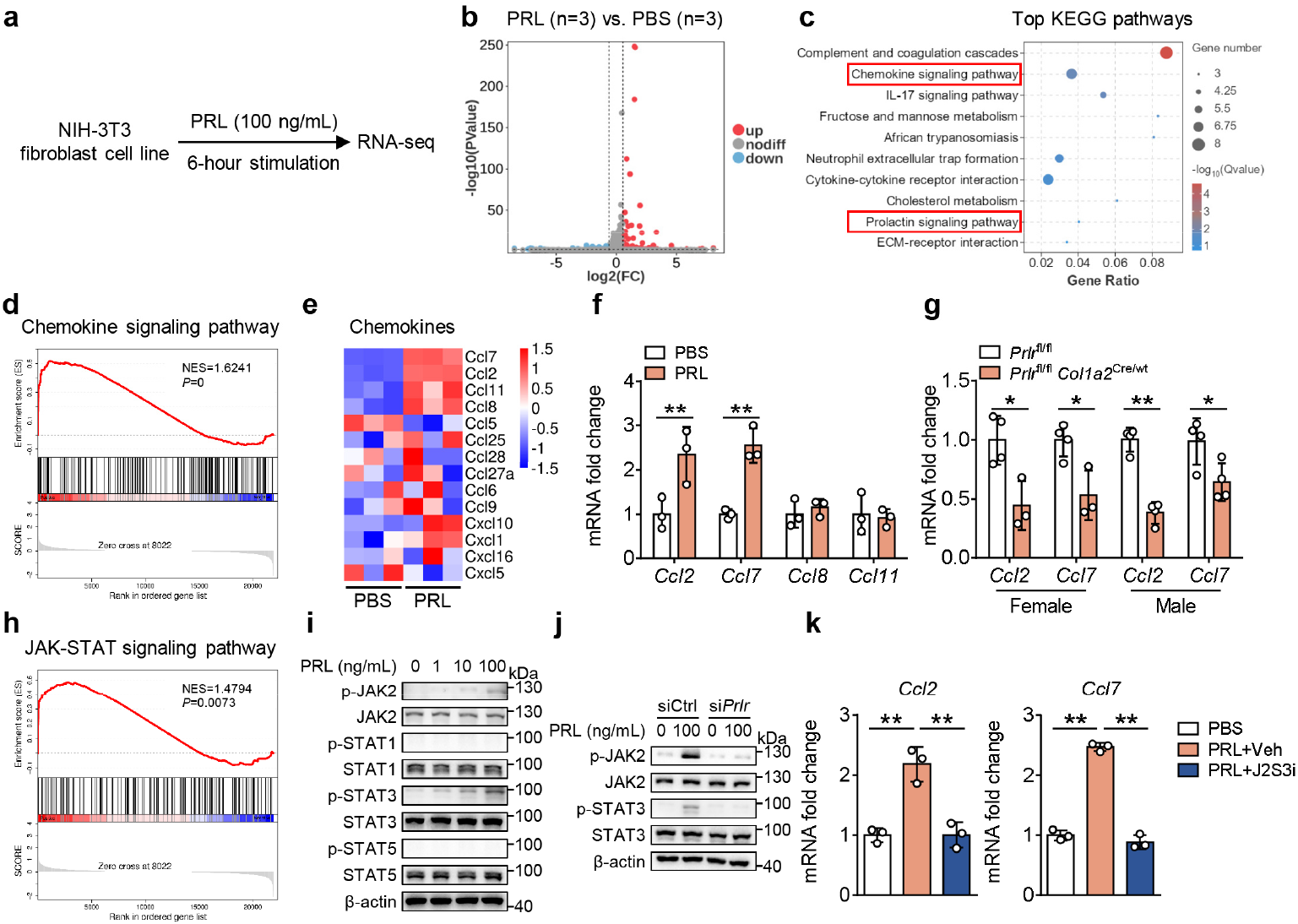
PRL promotes the production of CCL2 and CCL7 in fibroblasts via the JAK2-STAT3 signaling. **a** Schematic representation of the cell culture. **b** Volcano plot of differentially expressed genes between PBS-and PRL-stimulated NIH-3T3 cells (*n* = 3). **c** KEGG pathway enrichment analysis of differentially expressed genes. **d** GSEA gene enrichment analysis of the chemokine signaling pathway. **e** Heatmap of chemokines. **f** The expression of *Ccl2, Ccl7, Ccl8*, and *Ccl11* as determined by quantitative PCR (*n* = 3). **g** The expression of *Ccl2* and *Ccl7* in the skin of IMQ-treated female and male mice (*n* = 3–4). **h** GSEA gene enrichment analysis of the JAK-STAT signaling pathway. **i** Western blot analyses of p-JAK2, JAK2, p-STAT1, STAT1, p-STAT3, STAT3, p-STAT5, and STAT5 in PRL-stimulated NIH-3T3 cells. **j** Western blot analyses of p-JAK2, JAK2, p-STAT3, and STAT3 as indicated. **k** The expression of *Ccl2* and *Ccl7* in PBS-, PRL+Veh-, and PRL+J2S3i-treated NIH-3T3 cells (*n* = 3). Data are representative of two (**f, g, i**–**k**) independent experiments. Data are mean ± SD (**f, g, k**). The *P* values were calculated by two-tailed unpaired Student’s *t*-test (**f, g**) or one-way ANOVA with Dunnett’s post-hoc test (**k**); \**P* < 0.05, \*\**P* < 0.01.

We then confirmed the role of the PRL-JAK2-STAT3-CCL2/CCL7 axis in IMQ-induced psoriasis *in vivo*. PRL-aggravated psoriasis was alleviated in CCR2 (the receptor of CCL2 and CCL7, highly expressed in monocytes/macrophages^26^) inhibitor- or CCL2/CCL7 inhibitor-administered mice relative to vehicle-administered mice, as evidenced by a decrease in neutrophils, monocytes, macrophages, and IL-17A^+^ γδTCR^lo^ cells in the skin (**Fig. 6**). Similarly, a JAK2/STAT3 inhibitor administration alleviated PRL-aggravated psoriasis and reduced the expression of *Ccl2, Ccl7, Tnf, Il1b, Il17a*, and *Il23a* in the skin (**Supplementary Fig. 8**). Altogether, these data indicate that PRL-PRLR signaling in fibroblasts promotes the production of CCL2 and CCL7 via the JAK2-STAT3 signaling. These chemokines orchestrate a pathogenic immune cell response that amplifies psoriasiform dermatitis.

**Fig. 6.**
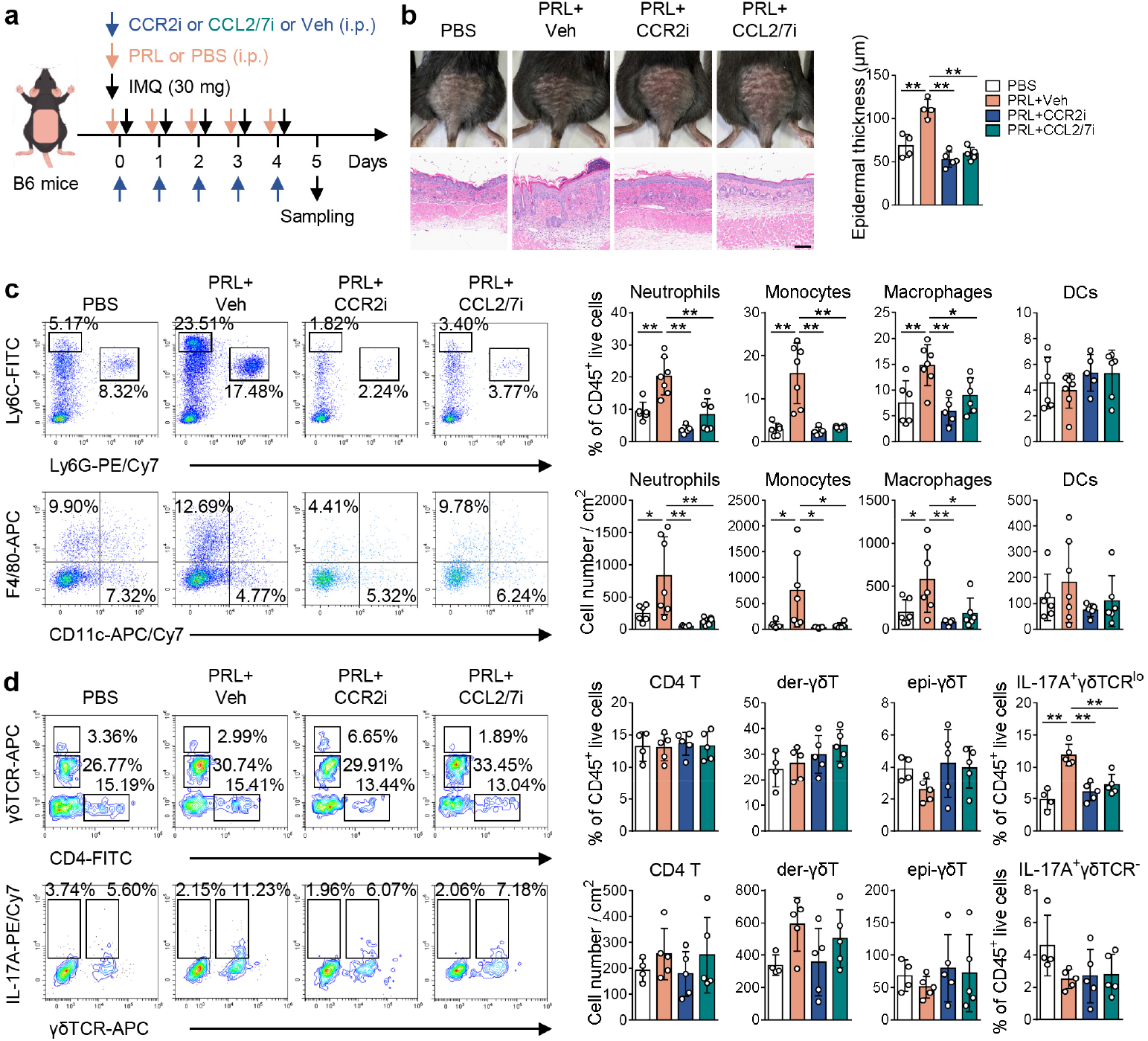
Blocking the CCL2/CCL7-CCR2 signaling ameliorates PRL-aggravated psoriasis. **a** Schematic representation of the mouse model. **b** Representative photos of mouse back skin and H&E staining images, and statistical analysis of epidermal thickness in PBS, PRL+Veh, PRL+CCR2i, and PRL+CCL2/7i mice (*n* = 4–5). Scale bar, 100 μm. **c, d** Representative flow cytometry plots and statistical analyses of neutrophils, monocytes, macrophages, and DCs in the skin (*n* = 5–7) (**c**) and CD4 T, der-γδT, epi-γδT, IL-17A^+^ γδTCR^lo^, and IL-17A^+^ γδTCR^-^ cells in the skin (*n* = 4–5) (**d**). Numbers in representative plots indicate the frequency of each cell subset among CD45^+^ cells. Data are from two independent experiments. Data are mean ± SD (**b**–**d**). The *P* values were calculated by one-way ANOVA with Dunnett’s post-hoc test (**b**–**d**); \**P* < 0.05, \*\**P* < 0.01.

### Pharmacological targeting of PRL signaling alleviates psoriasis in mice and human skin explants

We next assessed the possibility of targeting PRL in psoriasis therapy. IMQ-induced psoriasis mice treated with PRL inhibitor bromocriptine (BRC, inhibiting the production of the pituitary PRL) or PRL neutralizing antibody anti-PRL displayed less severe IMQ-induced psoriasiform dermatitis compared with the control mice (**Fig. 7**). The observed alleviation coincided with reduced serum PRL levels, downregulated expression of *Ccl2* and *Ccl7*, and decreased numbers of neutrophils, monocytes, macrophages, and IL-17A^+^ γδTCR^lo^ cells in the skin (**Fig. 7**).

**Fig. 7.**
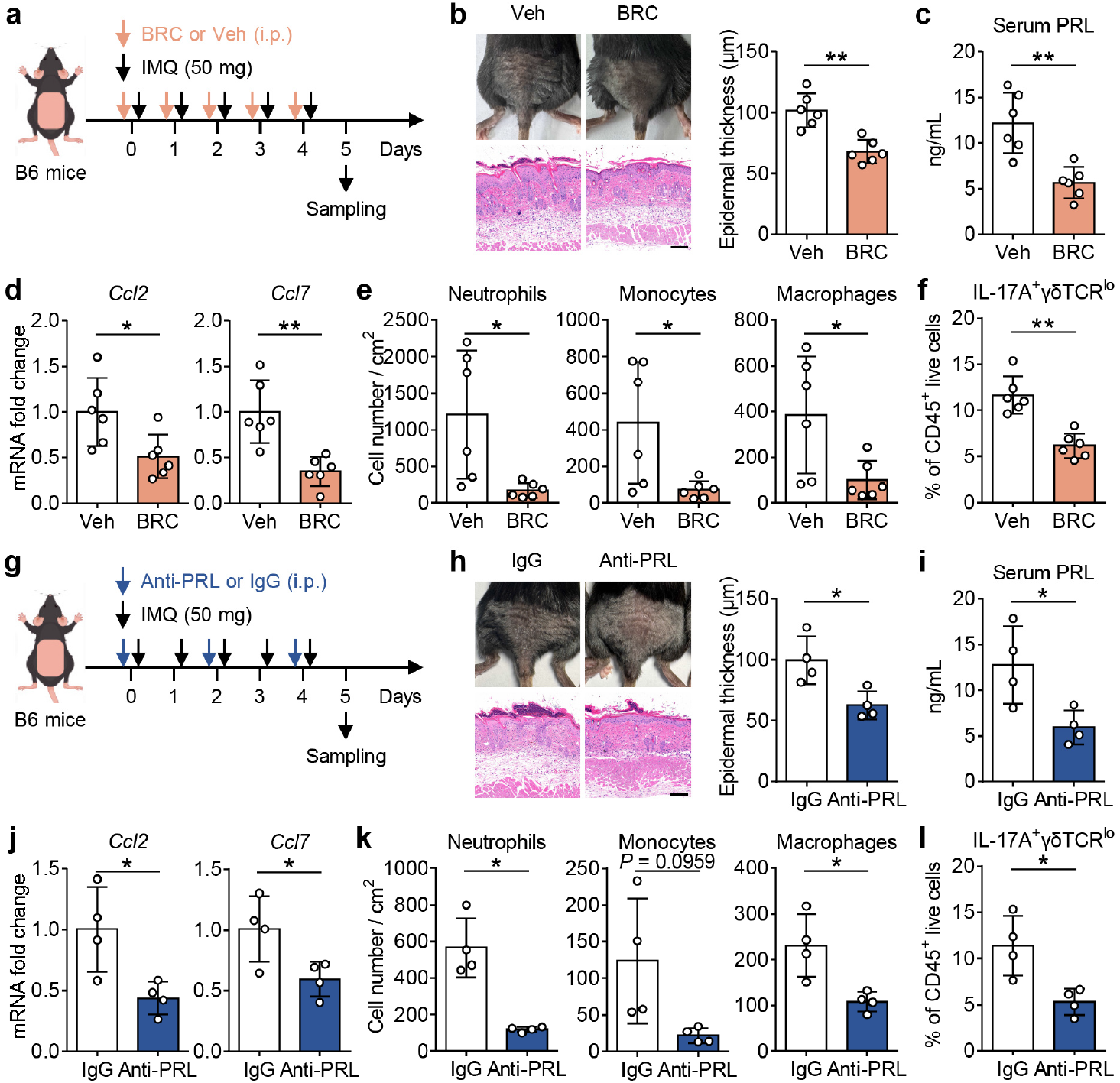
Pharmacological targeting of PRL alleviates psoriasis. **a** Schematic representation of the mouse model. **b**–**f** Representative photos of mouse back skin and H&E staining images, and statistical analysis of epidermal thickness (**b**), serum PRL levels (**c**), the expression of *Ccl2* and *Ccl7* (**d**), the numbers of neutrophils, monocytes, and macrophages (**e**), and the percentage of IL-17A^+^ γδTCR^lo^ cells (**f**) in Veh and BRC mice (*n* = 6). **g** Schematic representation of the mouse model. **h**–**l** Representative photos of mouse back skin and H&E staining images, and statistical analysis of epidermal thickness (**h**), serum PRL levels (**i**), the expression of *Ccl2* and *Ccl7* (**j**), the numbers of neutrophils, monocytes, and macrophages (**k**), and the percentage of IL-17A^+^ γδTCR^lo^ cells (**l**) in IgG and anti-PRL mice (*n* = 4). **b, h** Scale bar, 100 μm. Data are from one (**g**–**l**) or two (**a**–**f**) independent experiments. Data are mean ± SD (**b**–**f, h**–**l**). The *P* values were calculated by two-tailed unpaired Student’s (**b**–**d, f, h**–**j, l**) or Welch’s (**e, k**) *t*-test; \**P* < 0.05, \*\**P* < 0.01.

Finally, we investigated the clinical relevance between PRL and psoriasis. We found that serum PRL levels were positively correlated with PASI as well as serum levels of TNF-α, IL-17A, CCL2, and CCL7 (**Fig. 8a**). Next, we modeled psoriasiform dermatitis in normal human skin by directly stimulating skin explants with TNF-α, IFN-γ, and IL-17A. In this *ex vivo* model, PRL-induced *CCL*2 and *CCL7* expression was suppressed by a JAK2/STAT3 inhibitor (**Fig. 8b**). The transwell chemotaxis assay further confirmed that the PRL-JAK2-STAT3-CCL2/CCL7 axis promotes the recruitment of monocytes (**Fig. 8c**). These data suggest that PRL signaling contributes to recruiting monocytes into skin in human psoriasis.

**Fig. 8.**
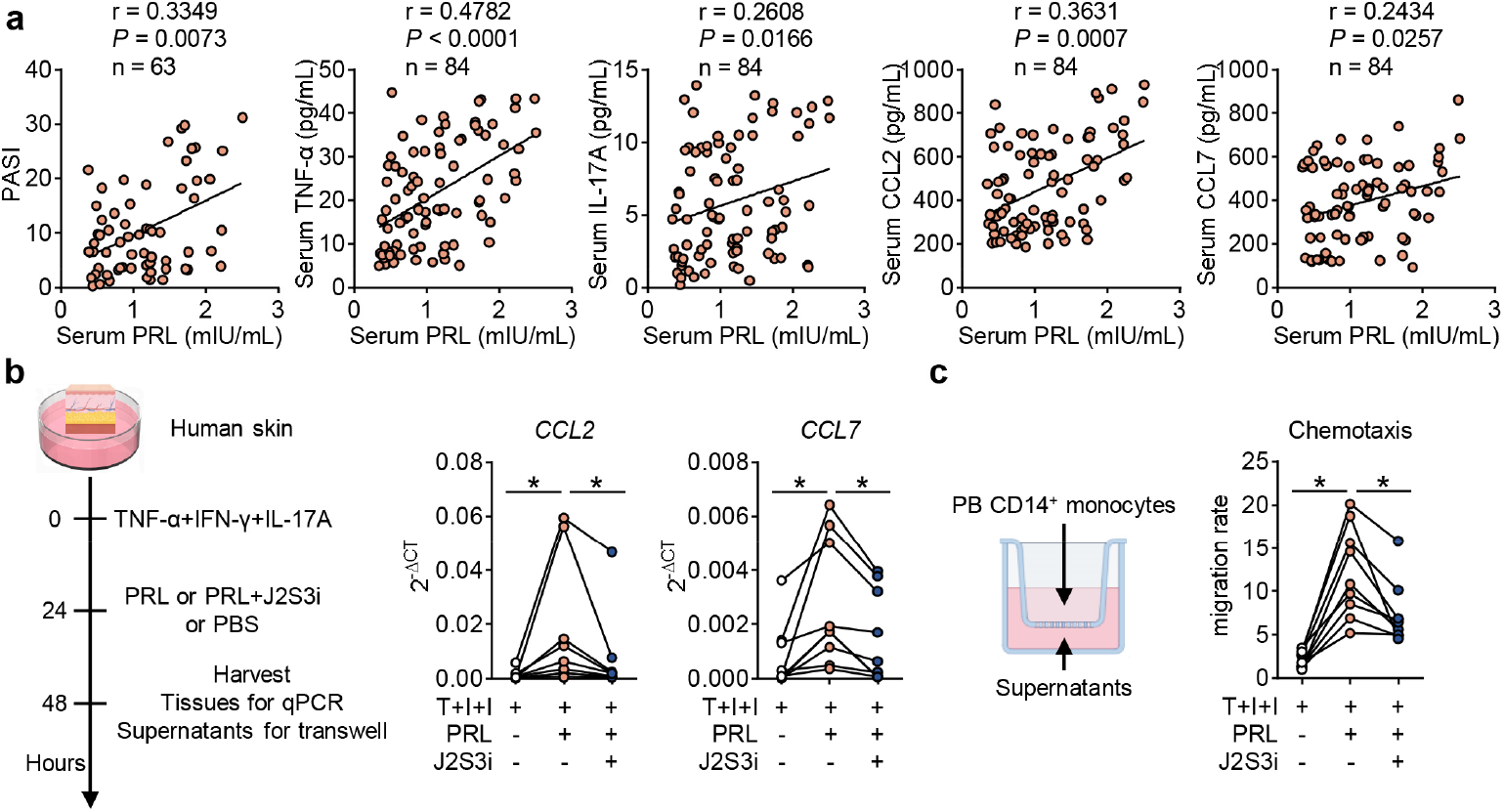
Clinical relevance between PRL and psoriasis. **a** Correlations of serum PRL levels with PASI (*n* = 63), as well as with serum levels of TNF-α, IL-17A, CCL2, and CCL7 (*n* = 84) in psoriasis patients. **b** Schematic representation of the *ex vivo* human skin explant culture and the expression of *CCL2* and *CCL7* in human skin (*n* = 9). **c** Schematic representation of the transwell chemotaxis assay and statistical analysis of migration rate (*n* = 9). The *P* values were calculated by Spearman’s rank correlation test (**a**) or Wilcoxon signed-rank test (**b, c**); \**P* < 0.05.

## Discussion

The findings of this study clearly show a mechanism linking psychological stress to the pathogenesis of psoriasis. The HP axis-derived hormone PRL promotes local skin immune inflammation by acting on PRLR in fibroblasts, thereby playing a critical role in psoriasis pathology (**Supplementary Fig. 9**).

We first confirmed a positive correlation between psoriasis and psychological stress at both clinical and animal model levels, which is consistent with a number of clinical observations and experimental studies^27-30^. Subsequently, we found that the HP axis was activated in a mouse model of psoriasis, with elevated levels of multiple pituitary hormones, among which PRL attracted our particular attention. This is because analysis of single-cell RNA sequencing data identified that the expression of PRLR was specifically expressed in skin cells, being high in fibroblasts but low in KCs. This finding was validated in both human and mouse skin samples. More importantly, psoriasis patients with anxiety and/or depression had higher serum PRL levels, suggesting that stress may exacerbate psoriasis by elevating PRL levels. Previous studies have attributed stress-induced exacerbation of psoriasis to the activation of the HPA axis and the release of dysfunctional glucocorticoids^9,31,32^, now our findings shift the focus to another pituitary hormone PRL.

To identify the pathogenic target cell of PRL, we generated mice with fibroblast- or KC-specific knockout of *Prlr*. Functional experiments confirmed that specific deletion of *Prlr* in fibroblasts significantly alleviated IMQ-induced psoriasiform dermatitis, whereas deletion in KCs had no such effect. Furthermore, exogenous PRL supplementation exacerbated the disease, but this effect was abolished in fibroblast-specific *Prlr* knockout mice. This evidence strongly indicates that PRL promotes psoriasis primarily via PRLR in fibroblasts. This finding extends beyond the previous view of PRL as a general pro-inflammatory factor by precisely identifying its cell-specific role in psoriasis^33,34^. Further investigation revealed that AR-induced stress could elevate PRL levels and worsen psoriasis, and deletion of *Prlr* in fibroblasts partially ameliorated this stress-aggravated phenotype, providing direct experimental evidence for stress exacerbating psoriasis through the PRL-PRLR^+^ fibroblasts axis.

Mechanistically, we found that in fibroblasts, PRL specifically induces the expression of CCL2 and CCL7 by activating the JAK2-STAT3 signaling, rather than STAT1 or STAT5. This presents an interesting contrast to the conventional understanding that PRL activates STAT5 in other tissues, revealing the tissue specificity of PRL signaling^35^. *In vivo*, administration of a CCR2 inhibitor, a CCL2/CCL7 inhibitor, or a JAK2/STAT3 inhibitor effectively reversed the aggravating effect of PRL on psoriasiform dermatitis and reduced the recruitment of monocytes/macrophages and the frequency of IL-17A-producing γδT cells in the skin. The PRL-JAK2-STAT3-CCL2/CCL7 axis we delineated provides a novel upstream mechanism for understanding the recruitment of monocytes/macrophages and the amplification of the IL-17 inflammatory circuit in psoriasis. In turn, heightened IL-17A promotes anxiogenic behaviors by increasing the excitability of IL-17RA-expressing basolateral amygdala neurons^29^, thereby forming a vicious cycle of stress and psoriasis.

The findings of this study have significant translational value. We confirmed that pharmacological targeting of PRL signaling effectively alleviated psoriasis in mice. More importantly, in *ex vivo* human skin explants, PRL similarly induced the expression of CCL2 and CCL7, a process that could be inhibited by a JAK2/STAT3 inhibitor, suggesting the PRL-JAK2-STAT3-CCL2/CCL7 axis is likely relevant in human psoriasis. Although JAK inhibitors have shown efficacy in treating psoriasis, their mechanisms are often attributed to the direct inhibition of signaling in immune cells^36,37^. Our results suggest that the partial efficacy of JAK inhibitors may be due to their inhibition of the PRL-fibroblast axis. Clinical correlation analysis showed that serum PRL levels in psoriasis patients positively correlated with disease severity and levels of inflammatory factors, further supporting the pathogenic role of PRL in human psoriasis. In conclusion, our study identifies PRL as a central molecular bridge connecting psychological stress and psoriasis and details the mechanism by which it drives immune inflammation via a fibroblast-chemokine pathway. These findings not only deepen our understanding of the “mind-skin” comorbidity in psoriasis but also provide a potential approach for treating psoriasis, particularly stress-associated psoriasis.

## Methods

### Mice

*Prlr*-flox (#NM-CKO-210028) and *Col1a2*-Cre (#NM-KI-215043) mice were purchased from Shanghai Model Organisms Center, Inc. (Shanghai, China). C57BL/6 (B6, Strain NO. N000013) and *KRT14*-Cre (Strain NO. T004833) mice were purchased from GemPharmatech (Nanjing, China). *Prlr*^fl/fl^ *Col1a2*^Cre/wt^ or *Prlr*^fl/fl^ *KRT14*^Cre/wt^ mice were generated by crossing *Prlr*-flox with *Col1a2*-Cre or *KRT14*-Cre mice, respectively. All mice were housed under a 12 h/12 h dark/light cycle, an ambient temperature of 22–25°C, and a humidity of 40–70% in specific-pathogen-free conditions at the Laboratory Animal Center of Ningbo University. Age- and sex-matched 8–10-week-old mice were used for experiments. Animal experiments were approved by the Animal Ethics and Welfare Committee of Ningbo University (Approval No.: NBU20240361) and conformed to the guidelines of the Guide for the Care and Use of Laboratory Animals. All efforts were made to minimize suffering.

### Human participants

Peripheral blood and skin samples were collected from psoriasis patients and healthy volunteers listed in Supplementary Table 1. Patients were diagnosed with psoriasis vulgaris by two senior dermatologists. Healthy volunteers had no diagnosed diseases at the time of collection. Written informed consent was obtained from all participants. Human sample studies were approved by the Medical Ethics Committee of Ningbo University (Approval No.: NBU-2024-084) and were conducted in accordance with the principles of the Declaration of Helsinki.

### Animal experiments

For establishment of psoriasis mouse model, mice were topically treated with 50 mg or 30 mg of 5% IMQ cream (MedShine) on shaved back skin (approximately 2 × 2 cm) daily for 5 consecutive days. For the establishment of stress mouse model, mice were restrained in a ventilated 50-mL tube for 4 h daily for 5 consecutive days. For exogenous PRL supplementation, mice were intraperitoneally injected with 4 μg of recombinant mouse PRL (MCE, #HY-P700261) daily for 5 consecutive days. For inhibiting CCL2/CCL7-CCR2 or JAK2/STAT3 signaling, mice were intraperitoneally injected with 0.2 mg of CCL2/CCL7 inhibitor AF2838 (MCE, #HY-B0498), 0.2 mg of CCR2 inhibitor INCB3344 (MCE, #HY-50674), or 0.2 mg of JAK2/STAT3 inhibitor FLLL32 (MCE, #HY-100544) daily for 5 consecutive days. For inhibiting PRL, mice were intraperitoneally injected with 0.2 mg of BRC (MCE, #HY-12705A) daily for 5 consecutive days or 30 μg of anti-PRL antibody (R&D, #AF1445) daily on days 0, 2, and 4.

### Cell culture

NIH-3T3 cells were cultured in DMEM containing 10% FBS and 1% penicillin-streptomycin at 37°C with 5% CO_2_. For RNA-seq, the cells were stimulated with recombinant human PRL (100 ng/mL, MCE, #HY-P71059) for 6 h. For Western blot, the cells were stimulated with different concentrations of PRL (0, 1, 10, and 100 ng/mL) for 30 min, or transfected with siRNA targeting *Prlr* (5’-UAAAGAAACAGGAAUUGGGTT-3’) or scrambled control for 24 h and then stimulated with PRL (100 ng/mL) for 30 min. For quantitative PCR, the cells were pretreated with JAK2/STAT3 inhibitor FLLL32 (10 μM) for 1 h followed by stimulation with PRL (100 ng/mL) for 6 h.

### *Ex vivo* human skin explant culture

Normal foreskin tissues (5 × 5 mm) were stimulated in DMEM supplemented with recombinant human TNF-α (10 ng/mL, ThermoFisher, #300-01A), IFN-γ (20 ng/mL, ThermoFisher, #300-02), and IL-17A (10 ng/mL, ThermoFisher, #200-17) for 24 h, and then stimulated with either recombinant human PRL (100 ng/mL) or JAK2/STAT3 inhibitor FLLL32 (10 μM) for another 24 h. After stimulation, skin tissues and culture supernatants were collected for subsequent experiments.

### Mouse behavioral test (elevated plus maze and open-field test)

Anxiety-like behavior was assessed in a 5-minute session on the elevated plus maze. Depression-like behavior was evaluated by allowing mice to freely explore a square arena (40 × 40 × 40 cm) for 10 minutes in the open-field test. Both tests were conducted during the light phase in a sound-attenuated room, with the experimenter blinded to the group allocation. All sessions were video-recorded and analyzed using an automated tracking system (VisuTrack v2.0).

### Single-cell RNA sequencing data analysis

The data downloaded from GSE165021 and GSE228421 were analyzed using the Seurat package (version 5.0.1) in R (version 4.4.1). Cells expressing 200–6,500 (in Fig. 2c) or 300–4,000 (in Fig. 2h) genes that were detected in at least 3 cells and a percentage of mitochondrial genes below 10% (in Fig. 2c) or 12% (in Fig. 2h) were considered viable and included for analysis. The data were then normalized and scaled. Dimensionality reduction was performed using *RunPCA*, followed by batch effect elimination using *RunHarmony*. Cells were clustered using *FindNeighbors* (a dimension of 25 (in Fig. 2c) or 30 (in Fig. 2h)) and *FindClusters* (a resolution of 0.2 (in Fig. 2c) or 0.5 (in Fig. 2h)).

### Cell isolation

To isolate skin immune cells, the back skin was separated, cut into small pieces, and incubated in DMEM (with 25 mM HEPES and 1.8 mM CaCl_2_) containing 2 mg/mL collagenase II (Sigma, #C6885), 50 μg/mL DNase I (Sangon Biotech, #A610099), and 10% FBS for 90 min (manually shake for 10 s every 30 min) in a 37°C shaking incubator (220 rpm). After adding additional DMEM with 10% FBS to inactivate enzyme activity, digested skin pieces were passed through 40-μm cell strainers. Skin leukocytes were isolated by 30% and 70% percoll (GE Healthcare, #17-0891-01) at 1260g for 20 min. The pellets were resuspended in PBS with 3% FBS, counted, and used for flow cytometry.

### Flow cytometry

Single-cell suspensions in PBS with 3% FBS were blocked with purified anti-mouse CD16/32 (BD Pharmingen, #553143, clone: 2.4G2) and then surfacely stained with fluorochrome-conjugated antibodies. Zombie Aqua dye (BioLegend, #423102) was used to identify live or dead cells. After washing with PBS with 3% FBS, the cells were fixed using 4% PFA and permeabilized using 0.5% Triton X-100, and intracellularly stained with fluorochrome-conjugated antibodies. All data were acquired and analyzed using a flow cytometer (Beckman Coulter CytoFLEX) with CytExpert software (version 2.4) and FlowJo software (version v10).

The following antibodies were used: PerCP/Cy5.5-CD45 (BioLegend, #103132, clone: 30-F11), FITC-Ly6C (BioLegend, #128006, clone: HK1.4), PE/Cy7-Ly6G (BioLegend, #127618, clone: 1A8), APC-F4/80 (BioLegend, #123116, clone: BM8), APC/Cy7-CD11c (BioLegend, #117324, clone: N418), FITC-CD4 (BioLegend, #100406, clone: GK1.5), APC-γδTCR (BioLegend, #118116, clone: GL3), and PE/Cy7-IL-17A (BioLegend, #506922, clone: TC11-18H10.1).

### RNA sequencing

Total RNA was extracted using Trizol reagent (ThermoFisher, #15596018CN) according to the manufacturer’s instructions. After quality control of RNA amount, purity, and integrity, mRNA was enriched by Oligo (dT) beads. Then the enriched mRNA was fragmented into short fragments (200–700nt) and reversely transcribed into cDNA. The cDNA library was sequenced using Illumina Novaseq 6000. Differentially expressed genes were defined as |log_2_FC| > log_2_1.5 and *P* < 0.05. Kyoto Encyclopedia of Genes and Genomes (KEGG) pathway enrichment analysis and Gene Set Enrichment Analysis (GSEA) were done. All services were provided by Gene Denovo Biotechnology Corporation (Guangzhou, China).

### RNA isolation, reverse transcription, and quantitative PCR

Total RNA was isolated using Trizol reagent according to the manufacturer’s instructions and then reverse-transcribed to cDNA using an *Evo M-MLV* RT Mix Kit with gDNA Clean for qPCR (Accurate Biology, #AG11728). Quantitative PCR was performed by a SYBR Green Premix *Pro Taq* HS qPCR Kit (Accurate Biology, #AG11701) with a Real-Time System (Roche LightCycler 480). Gene expression was calculated using the 2^−ΔΔCT^ method relative to housekeeping gene *Gapdh* or *GAPDH*. The sequences of primers were listed in Supplementary Table 2.

### Western blot

The cells were lysed in RIPA buffer (Beyotime, #P0013C) containing PMSF (Beyotime, #ST506) and protease and phosphatase inhibitor cocktail (Beyotime, #P1008). Western blot was performed according to standard protocols. The following antibodies were used: p-JAK2 (Abcam, #ab32101, 1:1,000), JAK2 (CST, #3230S, 1:1,000), p-STAT1 (CST, #9167S, 1:1,000), STAT1 (CST, #14994S, 1:1,000), p-STAT3 (CST, #9145S, 1:1,000), STAT3 (CST, #4904S, 1:1,000), p-STAT5 (CST, #9359S, 1:1,000), STAT5 (CST, #94205S, 1:1,000), β-actin (CST, #4970S, 1:1,000), and HRP-conjugated goat anti-rabbit IgG (H+L) (ZSGB-BIO, #ZB-2301, 1:20,000). The signals were detected by a SuperSignal West Femto Maximum Sensitivity Substrate Kit (ThermoFisher, #34095) and imaged with an imaging system (Tanon 5200).

### ELISA

Mouse PRL (mlbio, #ml001906), TSH (mlbio, #ml001956), FSH (mlbio, #ml001910), LH (mlbio, #ml001984), α-MSH (mlbio, #ml037968), β-EP (mlbio, #ml001926), ACTH (mlbio, #ml001895), and human PRL (mlbio, #ml058210) were measured using commercial ELISA kits according to the manufacturer’s instructions.

### Histological analysis

Skin tissues were fixed in 10% formalin, embedded in paraffin, and cut into 4-μm sections for hematoxylin and eosin (H&E) staining. The images were obtained using a microscope (Olympus BX53) with cellSens software (version 1.5). The epidermal thickness was calculated as the mean of three randomly selected areas by two blinded observers.

For immunofluorescence staining, deparaffinized sections were boiled for 15 min in 10 mM sodium citrate buffer (pH 6.0) for antigen retrieval. After blocking nonspecific binding, the sections were incubated with mouse anti-COL1A2 (Santa Cruze, #sc-393573, 1:50), mouse anti-KRT14 (Santa Cruze, #sc-53253, 1:50), and rabbit anti-PRLR (Abcam, #ab214303, 1:200) primary antibodies at 4°C overnight. Secondary antibodies were used including FITC-conjugated goat anti-mouse IgG (H+L) (ZSGB-BIO, #ZF-0312, 1:100) and Rhodamine-conjugated goat anti-rabbit IgG (H+L) (ZSGB-BIO, #ZF-0316, 1:100). Nuclei were stained with DAPI (Beyotime, #P0131). The images were acquired using a confocal scanning microscope (Leica TCS SP8) with LAS X software (version 3.5.1).

### Transwell chemotaxis assay

Human peripheral monocytes from healthy controls were isolated by positive selection using PE-CD14 (BD Pharmingen, #555398, clone: M5E2) with anti-PE microbeads (Miltenyi Biotec, #130-048-801) and a MACS system. The purity of sorted cells was > 90%. For the chemotaxis assay, 1 × 10^4^ monocytes resuspended in serum-free medium were seeded into the upper chamber of a 24-well insert (5-μm pores). The lower chamber contained with culture supernatants from *ex vivo* human skin explant. After a 4-hour incubation at 37°C, the cells that had migrated to the lower chamber were counted. The migration rate was calculated as (number of migrated cells / total number of cells plated) × 100%.

### Statistical analysis

Data were analyzed using GraphPad Prism 8 software (version 8.0.1). After assessing normality via quantile-quantile plots and the Shapiro-Wilk test, group comparisons were performed as follows: for two independent groups, normally distributed data were analyzed using the two-tailed unpaired Student’s (equal variances, as assessed by *F*-test) or Welch’s (unequal variances) *t*-test, while non-normal data were analyzed using the Mann-Whitney U test; for two paired groups, the Wilcoxon signed-rank test was used; for multiple groups, one-way with Dunnett’s post-hoc test, or two-way ANOVA with Tukey’s post-hoc test was employed. Correlation was evaluated using the Spearman’s rank correlation test. *P* < 0.05 was considered statistically significant. Normally and non-normally distributed data are presented as mean ± standard deviation (SD) and median (interquartile range, IQR), respectively.

## Data availability

The data that support the findings of this study are available from the corresponding author upon reasonable request.

## Acknowledgements

This work was supported by National Natural Science Foundation of China (82404130 to Y.L.), Zhejiang Provincial Natural Science Foundation of China (LQ24H110001 to Y.L. and ZCLQN25H1101 to H.G.), and Ningbo Natural Science Foundation (2024J475 to Y.L.).

## Author contributions

L.Y., X.H., and D.F. conceived the experiments, supervised the project, and analyzed the data. H.G., S.Z., and Y.Z. designed and conducted the experiments and analyzed the data. X.Z. and M.C. fed and identified the mice. P.G., S.L., and Q.M. coordinated the clinical investigation. H.G., S.Z., and Y.Z. wrote the manuscript. All authors reviewed and revised the manuscript.

## References

1 Griffiths, C. E. M., Armstrong, A. W., Gudjonsson, J. E. & Barker, J. Psoriasis. Lancet 397, 1301–1315 (2021).

2 Parisi, R. et al. National, regional, and worldwide epidemiology of psoriasis: systematic analysis and modelling study. BMJ 369, m1590 (2020).

3 Torales, J. et al. Psychodermatological mechanisms of psoriasis. Dermatol Ther 33, e13827 (2020).

4 Rousset, L. & Halioua, B. Stress and psoriasis. Int J Dermatol 57, 1165–1172 (2018).

5 Ingram, J. R. & Ahluwalia, A. The pharmacology of itch. Br J Dermatol 184, e1–e2 (2021).

6 Sanders, K. M. & Akiyama, T. The vicious cycle of itch and anxiety. Neurosci Biobehav Rev 87, 17–26 (2018).

7 Stojanovich, L. & Marisavljevich, D. Stress as a trigger of autoimmune disease. Autoimmun Rev 7, 209–213 (2008).

8 Zhu, T. H. et al. The Role of the Nervous System in the Pathophysiology of Psoriasis: A Review of Cases of Psoriasis Remission or Improvement Following Denervation Injury. Am J Clin Dermatol 17, 257–263 (2016).

9 Marek-Jozefowicz, L. et al. The Brain-Skin Axis in Psoriasis-Psychological, Psychiatric, Hormonal, and Dermatological Aspects. Int J Mol Sci 23 (2022).

10 Weiglein, A., Gaffal, E. & Albrecht, A. Probing the Skin-Brain Axis: New Vistas Using Mouse Models. Int J Mol Sci 23 (2022).

11 Ostrowski, S. M., Belkadi, A., Loyd, C. M., Diaconu, D. & Ward, N. L. Cutaneous denervation of psoriasiform mouse skin improves acanthosis and inflammation in a sensory neuropeptide-dependent manner. J Invest Dermatol 131, 1530–1538 (2011).

12 Yin, Q. et al. Lidocaine Ameliorates Psoriasis by Obstructing Pathogenic CGRP Signaling-Mediated Sensory Neuron-Dendritic Cell Communication. J Invest Dermatol 142, 2173–2183 e2176 (2022).

13 Wang, Y. et al. Stress aggravates and prolongs imiquimod-induced psoriasis-like epidermal hyperplasis and IL-1beta/IL-23p40 production. J Leukoc Biol 108, 267–281 (2020).

14 Riol-Blanco, L. et al. Nociceptive sensory neurons drive interleukin-23-mediated psoriasiform skin inflammation. Nature 510, 157–161 (2014).

15 Cai, X. et al. Tenascin C(+) papillary fibroblasts facilitate neuro-immune interaction in a mouse model of psoriasis. Nat Commun 14, 2004 (2023).

16 Mbiydzenyuy, N. E. & Qulu, L. A. Stress, hypothalamic-pituitary-adrenal axis, hypothalamic-pituitary-gonadal axis, and aggression. Metab Brain Dis 39, 1613–1636 (2024).

17 Plikus, M. V. et al. Fibroblasts: Origins, definitions, and functions in health and disease. Cell 184, 3852–3872 (2021).

18 Korsunsky, I. et al. Cross-tissue, single-cell stromal atlas identifies shared pathological fibroblast phenotypes in four chronic inflammatory diseases. Med 3, 481–518 e414 (2022).

19 Xu, Z. et al. Anatomically distinct fibroblast subsets determine skin autoimmune patterns. Nature 601, 118–124 (2022).

20 Driskell, R. R. et al. Distinct fibroblast lineages determine dermal architecture in skin development and repair. Nature 504, 277–281 (2013).

21 Zhang, B. et al. Multitranscriptome analysis reveals stromal cells in the papillary dermis to promote angiogenesis in psoriasis vulgaris. Br J Dermatol 192, 672–683 (2025).

22 Ma, F. et al. Single cell and spatial sequencing define processes by which keratinocytes and fibroblasts amplify inflammatory responses in psoriasis. Nat Commun 14, 3455 (2023).

23 Li, Z. et al. Cross-disease characterization of fibroblast heterogeneities and their pathogenic roles in skin inflammation. Clin Immunol 255, 109742 (2023).

24 Qing, H. et al. Origin and Function of Stress-Induced IL-6 in Murine Models. Cell 182, 372–387 e314 (2020).

25 Chasseloup, F., Bernard, V. & Chanson, P. Prolactin: structure, receptors, and functions. Rev Endocr Metab Disord 25, 953–966 (2024).

26 Zlotnik, A. & Yoshie, O. The chemokine superfamily revisited. Immunity 36, 705–716 (2012).

27 Liu, S. et al. Triggers for the onset and recurrence of psoriasis: a review and update. Cell Commun Signal 22, 108 (2024).

28 Snast, I. et al. Psychological stress and psoriasis: a systematic review and meta-analysis. Br J Dermatol 178, 1044–1055 (2018).

29 Lee, B. et al. Inflammatory and anti-inflammatory cytokines bidirectionally modulate amygdala circuits regulating anxiety. Cell 188, 2190–2202 e2115 (2025).

30 Guo, J. et al. Depressive-like behaviors in mice with Imiquimod-induced psoriasis. Int Immunopharmacol 89, 107057 (2020).

31 Rajasekharan, A. et al. Stress and psoriasis: Exploring the link through the prism of hypothalamo-pituitary-adrenal axis and inflammation. J Psychosom Res 170, 111350 (2023).

32 Ge, H. et al. Stress aggravates imiquimod-induced psoriasiform inflammation by promoting M1 macrophage polarization. Int Immunopharmacol 124, 110899 (2023).

33 Hau, C. S. et al. Prolactin induces the production of Th17 and Th1 cytokines/chemokines in murine Imiquimod-induced psoriasiform skin. J Eur Acad Dermatol Venereol 28, 1370–1379 (2014).

34 Yang, H. et al. Local production of prolactin in lesions may play a pathogenic role in psoriatic patients and imiquimod-induced psoriasis-like mouse model. Exp Dermatol 27, 1245–1253 (2018).

35 Zhu, J. et al. Hepatic-derived extracellular vesicles in late pregnancy promote mammary gland development by stimulating prolactin receptor-mediated JAK2/STAT5/mTOR signalling. Int J Biol Macromol 281, 136498 (2024).

36 Bonelli, M. et al. Selectivity, efficacy and safety of JAKinibs: new evidence for a still evolving story. Ann Rheum Dis 83, 139–160 (2024).

37 Yu, J. et al. Pathogenesis, multi-omics research, and clinical treatment of psoriasis. J Autoimmun 133, 102916 (2022).

