## Supplementary Information for "Prolactin links psychological stress to psoriasis via a fibroblast-chemokine pathway"

**This file includes:**

Supplementary Figs. 1–9

Supplementary Tables 1 and 2

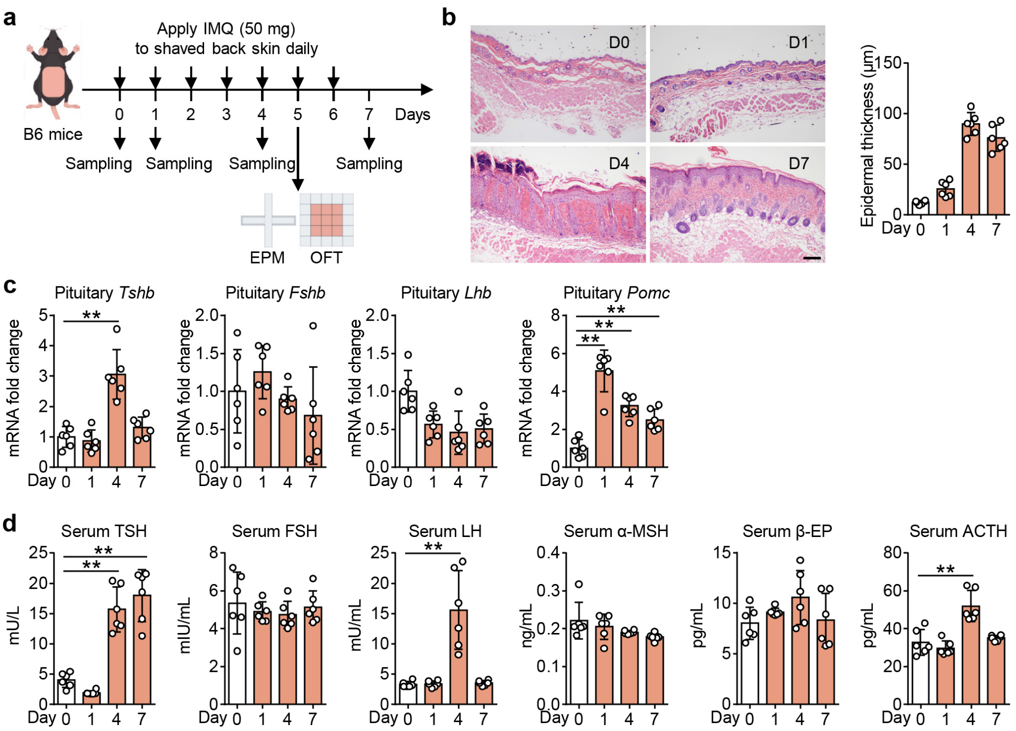


**Supplementary Fig. 1. Pituitary mRNA levels and serum protein levels of pituitary-secreted hormones in IMQ-induced psoriasis mice.**

**a** Schematic representation of the mouse model.

**b** Representative H&E staining images and statistical analysis of epidermal thickness in IMQ-treated mice at the indicated timepoints ($n$ = 6). Scale bar, 100 μm.

**c** Pituitary mRNA levels of thyroid stimulating hormone subunit beta (*Tshb*), follicle stimulating hormone subunit beta (*Fshb*), luteinizing hormone subunit beta (*Lhb*), and proopiomelanocortin (*Pomc*) in IMQ-treated mice at the indicated timepoints ($n$ = 6).

**d** Serum protein levels of thyroid stimulating hormone (TSH), follicle stimulating hormone (FSH), luteinizing hormone (LH), α-melanocyte stimulating hormone (α-MSH), β-endorphin (β-EP), and adrenocorticotropic hormone (ACTH) in IMQ-treated mice at the indicated timepoints ($n$ = 6).

Data are from two independent experiments. Data are mean ± SD (**b**–**d**). The $P$ values were calculated by one-way ANOVA with Dunnett's post-hoc test (**c**, **d**); $**P < 0.01$.

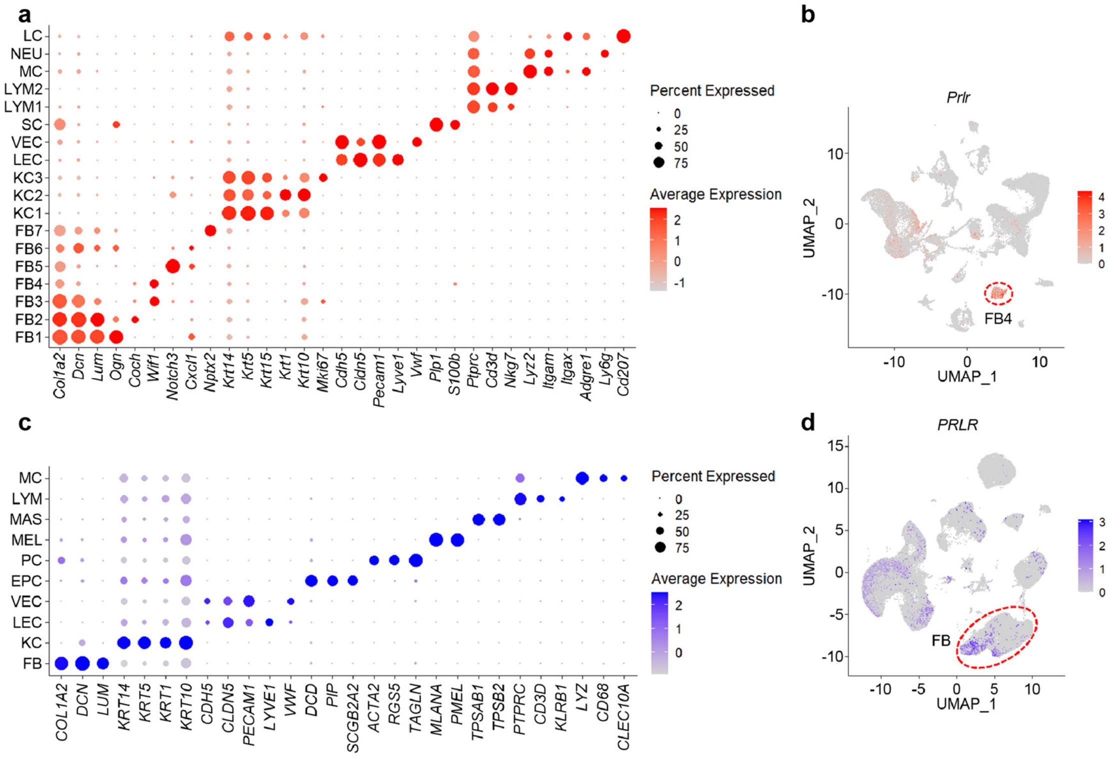


**Supplementary Fig. 2. Single-cell RNA sequencing data analysis of mouse and human skin.**

**a** Dot plot showing the expression of marker genes for annotating cell identities in mouse skin.

**b** Feature plot of *Prlr* expression.

**c** Dot plot showing the expression of marker genes for annotating cell identities in human skin.

**d** Feature plot of *PRLR* expression.

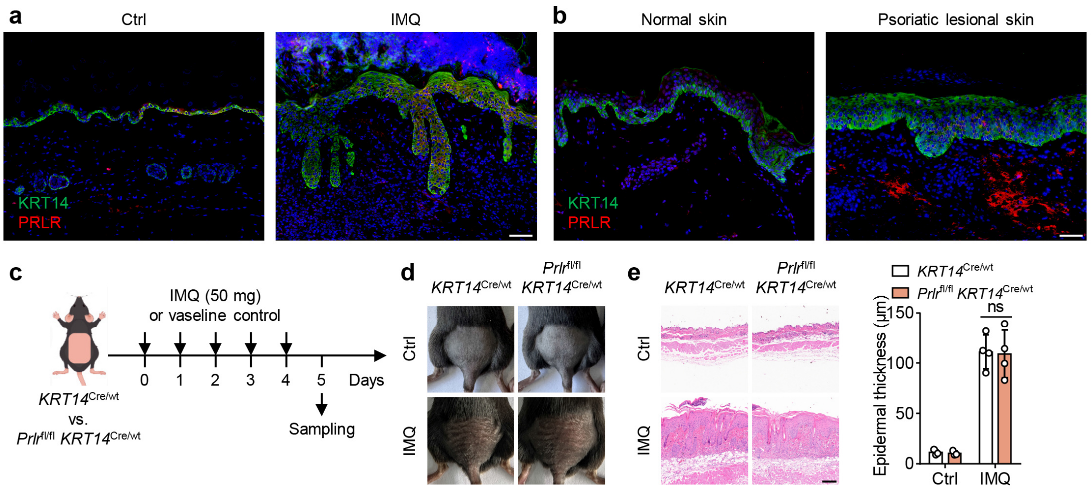


**Supplementary Fig. 3. Deletion of *Prlr* in KCs has no effect on IMQ-induced psoriasis.**

**a**, **b** Representative immunofluorescent images of skin sections stained with anti-KRT14 (green), anti-PRL (red), and DAPI (blue) in mouse ($n$ = 6) (**a**) and human ($n$ = 4) samples (**b**). The solid arrows show PRLR$^+$ KCs. Scale bar, 20 μm.

**c** Schematic representation of the mouse model.

**d** Representative photos of mouse back skin.

**e** Representative H&E staining images and statistical analysis of epidermal thickness in Ctrl and IMQ mice ($n$ = 3–4). Scale bar, 100 μm.

Data are from one (**c**–**e**) independent experiment. Data are mean ± SD (**e**). The $P$ values were calculated by two-way ANOVA with Tukey's post-hoc test (**e**); ns, not significant.

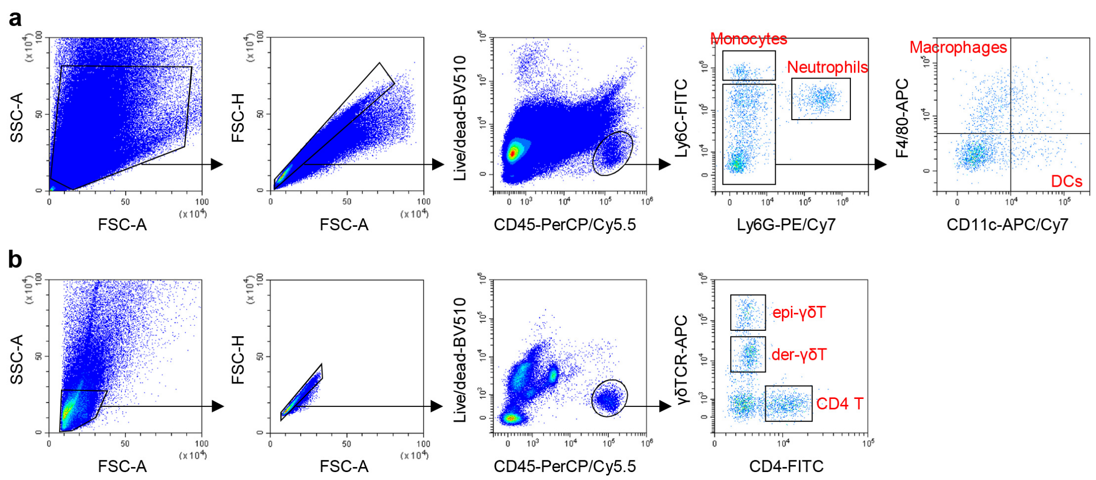


**Supplementary Fig. 4. The gating strategy for the identification of mouse skin myeloid cell and T cell subsets.**

**a**, **b** Representative flow cytometry plots showing gating strategy for identifying myeloid cell (**a**) and T cell (**b**) subsets in mouse skin.

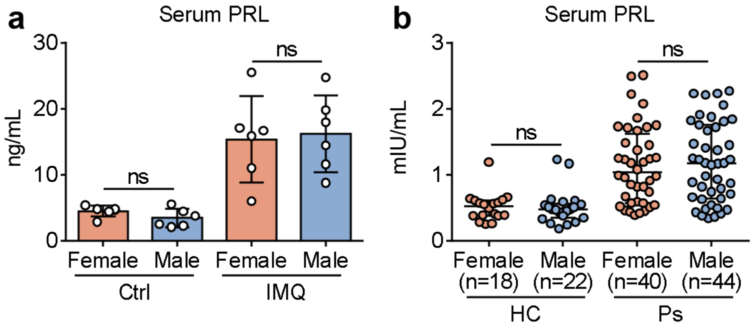


**Supplementary Fig. 5. Comparison of serum PRL levels between females and males in both mice and humans.**

**a** Serum PRL levels of female and male mice (*n* = 6).

**b** Serum PRL levels of female (*n* = 18) and male (*n* = 22) healthy controls, and female (*n* = 40) and male (*n* = 44) psoriasis patients.

Data are from two (**a**) independent experiments. Data are mean ± SD (**a**) or median (IQR) (**b**). The *P* values were calculated by two-tailed unpaired Student's *t*-test (**a**) or Mann-Whitney U test (**b**); ns, not significant.

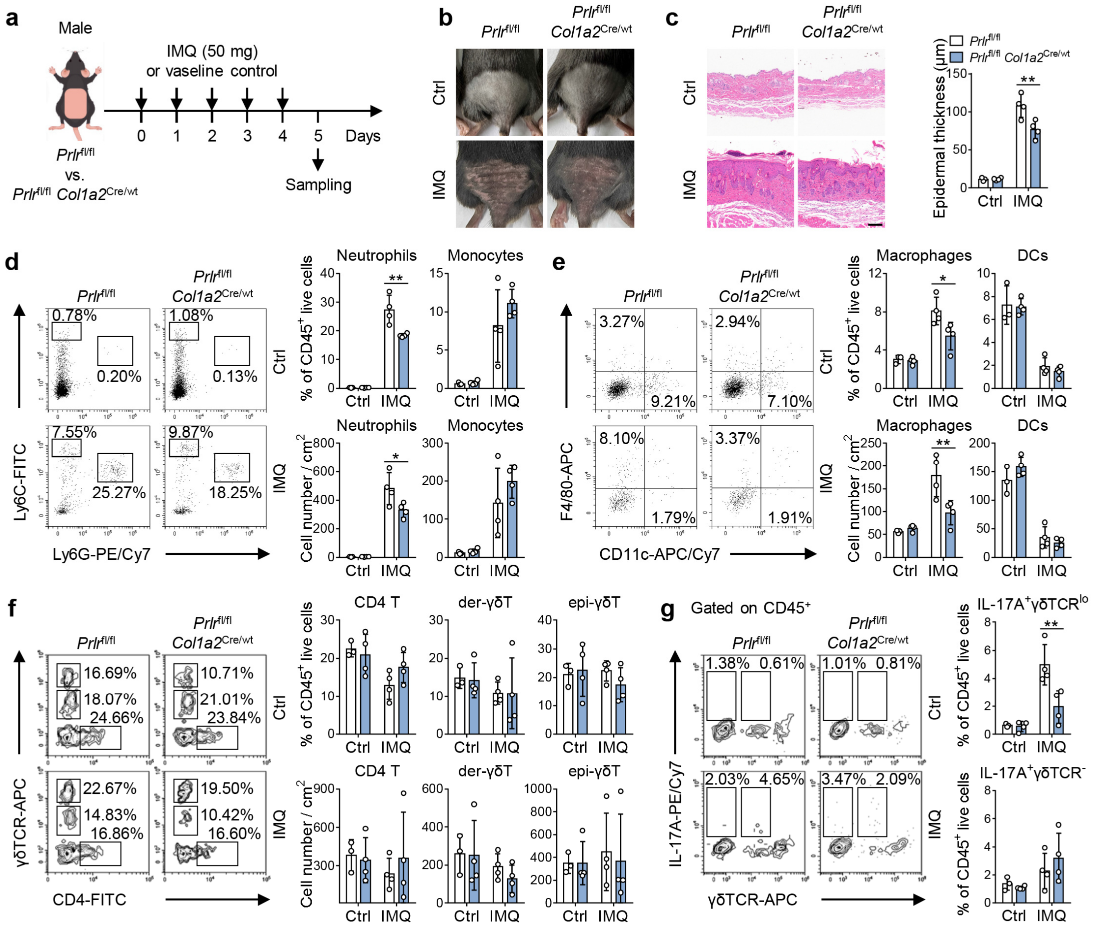


**Supplementary Fig. 6. Deletion of *Prlr* in fibroblasts alleviates IMQ-induced psoriasis in male mice.**

**a** Schematic representation of the mouse model.

**b** Representative photos of mouse back skin.

**c** Representative H&E staining images and statistical analysis of epidermal thickness in Ctrl and IMQ mice ($n$ = 3–4). Scale bar, 100 μm.

**d**–**g** Representative flow cytometry plots and statistical analyses of neutrophils and monocytes (**d**), macrophages and DCs (**e**), CD4 T, der-γδT, and epi-γδT cells (**f**), and IL-17A$^{+}$ γδTCR$^{lo}$ and IL-17A$^{+}$ γδTCR$^{-}$ cells (**g**) in the skin of Ctrl and IMQ mice ($n$ = 3–4). Numbers in representative plots indicate the frequency of each cell subset among CD45$^{+}$ cells.

Data are from one independent experiment. Data are mean ± SD (**c**–**g**). The $P$ values were calculated by two-way ANOVA with Tukey's post-hoc test (**c**–**g**); $*P < 0.05$, $**P < 0.01$.

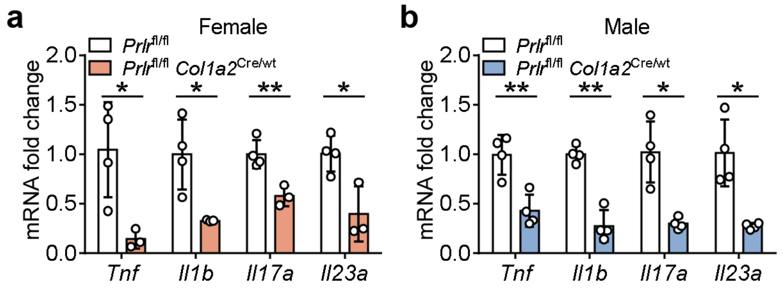


**Supplementary Fig. 7. Detection of pathogenic cytokines in the skin of female and male IMQ-treated mice.**

**a**, **b** The expression of pathogenic cytokines in the skin of IMQ-treated female (**a**) and male (**b**) mice (*n* = 3–4).

Data are from one independent experiment. Data are mean ± SD. The *P* values were calculated by two-tailed unpaired Student's or Welch's *t*-test; $^{*}P < 0.05$, $^{**}P < 0.01$.

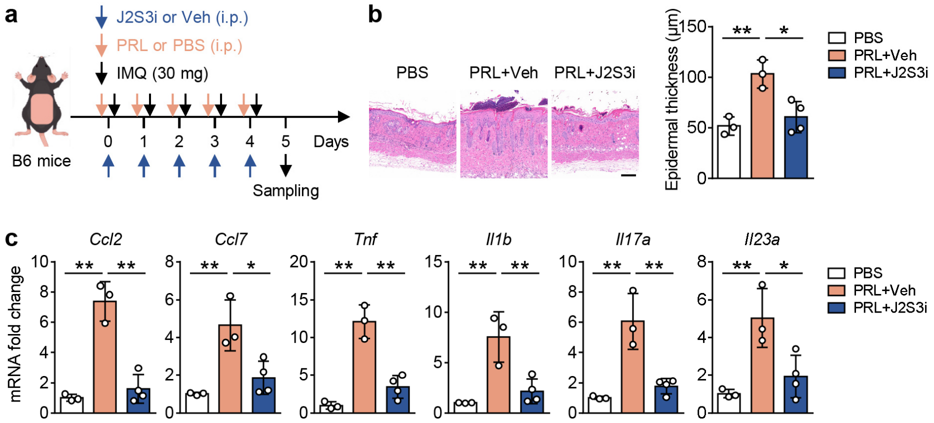


**Supplementary Fig. 8. Inhibition of JAK2/STAT3 signaling alleviates PRL-aggravated psoriasis.**

**a** Schematic representation of the mouse model.

**b** Representative H&E staining images and statistical analysis of epidermal thickness in PBS, PRL+Veh, and PRL+J2S3i mice (*n* = 3–4). Scale bar, 100 µm.

**c** The expression of pathogenic factors in the skin of PBS, PRL+Veh, and PRL+J2S3i mice (*n* = 3–4).

Data are from one independent experiment. Data are mean ± SD (**b**, **c**). The *P* values were calculated by one-way ANOVA with Dunnett's post-hoc test (**b**, **c**); $*P < 0.05$, $**P < 0.01$.

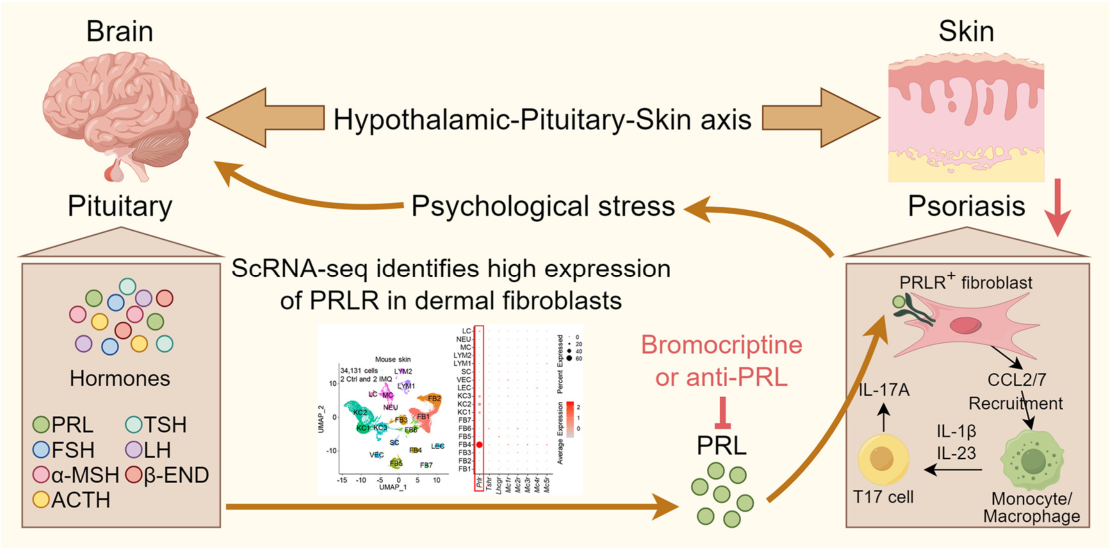


**Supplementary Fig. 9. Graphical summary.**

Psychological stress-induced PRL acts on dermal PRLR-expressing fibroblasts to promote the production of CCL2 and CCL7. These chemokines then recruit monocytes/macrophages into psoriatic lesional skin, thereby activating IL-17A-producing T cells through release IL-1β and IL-23. IL-17A not only fuels the development of psoriasis but also triggers anxiogenic behaviors, ultimately forming a vicious cycle of stress and psoriasis. Pharmacological targeting of PRL signaling alleviates psoriasis.

**Supplementary Table 1. Characteristics of healthy controls and psoriasis patients**

| Group | healthy controls | psoriasis patients |
|---|---|---|
| Total | 40 | 84 |
| Gender (female/male) | 18/22 | 40/44 |
| Age | 43 (26–63) | 46 (25–71) |
| PASI | / | 10.6 (0.3–31.2) |

**Supplementary Table 2. The sequences of primers**

| Genes | Forward primer (5’–3’) | Reverse primer (5’–3’) |
|---|---|---|
| *Gapdh* | GTGTTCCTACCCCCAATGTG | GGTCCTCAGTGTAGCCCAAG |
| *Ccl2* | CCAGCAAGATGATCCCAATG | TACGGGTCAACTTCACATTC |
| *Ccl7* | GCTGCTTTCAGCATCCAAGTG | CCAGGGACACCGACTACTG |
| *Ccl8* | CTGGGCCAGATAAGGCTCC | CATGGGGCACTGGATATTGTT |
| *Ccl11* | GAATCACCAACAACAGATGCAC | ATCCTGGACCCACTTCTTCTT |
| *Tnf* | ACTGGCAGAAGAGGCACTC | CTGGCACCACTAGTTGGTTG |
| *Il1b* | CTGAACTCAACTGTGAAATGC | TGATGTGCTGCTGCGAGA |
| *Il17a* | CTCAAAGCTCAGCGTGTCCAAACA | TATCAGGGTCTTCATTGCGGTGGA |
| *Il23a* | ATGCTGGATTGCAGAGCAGTA | ACGGGGCACATTATTTTTAGTCT |
| *GAPDH* | GTCTCCTCTGACTTCAACAGCG | ACCACCCTGTTGCTGTAGCCAA |
| *CCL2* | AGAATCACCAGCAGCAAGTGTCC | TCCTGAACCCACTTCTGCTTGG |
| *CCL7* | CCCTCACCCTCCAACATGAAA | TAGCTCTCCAGCCTCTGCTTA |
